# Magnetosomes could be protective shields against metal stress in magnetotactic bacteria

**DOI:** 10.1101/2020.06.02.128942

**Authors:** D. Muñoz, L. Marcano, R. Martín-Rodríguez, L. Simonelli, A. Serrano, A. García-Prieto, M.L. Fdez-Gubieda, A. Muela

**Affiliations:** Dpto. de Inmunología, Microbiología y Parasitología, Universidad del País Vasco - UPV/EHU, 48940 Leioa, Spain; Dpto. de Electricidad y Electrónica, Universidad del País Vasco - UPV/EHU, 48940 Leioa, Spain; Helmholtz-Zentrum Berlin für Materialen und Energie, Berlin, Germany; QUIPRE Department, University of Cantabria, 39005 Santander, Spain; Nanomedicine Group, IDIVAL, 39011 Santander, Spain; CLAESS beamline, ALBA Synchrotron, 08290 Cerdanyola del Vallès, Spain; SpLine, Spanish CRG BM25 Beamline, ESRF, 38000 Grenoble, France; Dpto. de Física Aplicada I, Universidad del País Vasco - UPV/EHU, 48013 Bilbao, Spain; BCMaterials, UPV/EHU Science Park, 48940 Leioa, Spain

**Keywords:** Magnetotactic bacteria, metal stress, magnetosomes, XANES, magnetite

## Abstract

Magnetotactic bacteria are aquatic microorganisms with the ability to biomineralise membrane-enclosed magnetic nanoparticles, called magnetosomes. These magnetosomes are arranged into a chain that behaves as a magnetic compass, allowing the bacteria to align in and navigate along the Earth’s magnetic field lines. According to the magneto-aerotactic hypothesis, the purpose of producing magnetosomes is to provide the bacteria with a more efficient movement within the stratified water column, in search of the optimal positions that satisfy their nutritional requirements. However, magnetosomes could have other physiological roles, as proposed in this work. Here we analyse the role of magnetosomes in the tolerance of *Magnetospirillum gryphiswaldense* MSR-1 to transition metals (Co, Mn, Ni, Zn, Cu). By exposing bacterial populations with and without magnetosomes to increasing concentrations of metals in the growth medium, we observe that the tolerance is significantly higher when bacteria have magnetosomes. The resistance mechanisms triggered in magnetosome-bearing bacteria under metal stress have been investigated by means of x-ray absorption near edge spectroscopy (XANES). XANES experiments were performed both on magnetosomes isolated from the bacteria and on the whole bacteria, aimed to assess whether bacteria use magnetosomes as metal storages, or whether they incorporate the excess metal in other cell compartments. Our findings reveal that the tolerance mechanisms are metal-specific: Mn, Zn and Cu are incorporated in both the magnetosomes and other cell compartments; Co is only incorporated in the magnetosomes, and Ni is incorporated in other cell compartments. In the case of Co, Zn and Mn, the metal is integrated in the magnetosome magnetite mineral core.

---

Magnetotactic bacteria (MTB) are aquatic microorganisms able to passively align parallel to the Earth’s geomagnetic field lines while they actively swim. This behavior, known as magnetotaxis, is due to the presence of unique intracellular magnetic organelles called magnetosomes^1–4^. The magnetosomes are intracellular inclusions composed by a core of magnetic iron mineral, typically magnetite (Fe_3_O_4_) or greigite (Fe_3_S_4_), enclosed by a thin membrane. Magnetotactic bacteria are widespread in freshwater and marine environments^5^. They are easily detected in chemically and redox stratified sediments and water columns, predominantly at the oxic-anoxic transition zones (OATZ). Bacteria living in OATZ, with vertical chemical gradients, are continually searching the optimal position in the stratified water column in order to satisfy their nutritional requirements. Under these circumstances, magnetotaxis is thought to be a great advantage by increasing the efficiency of chemotaxis^1^. Due to their inclination, the geomagnetic field lines act as vertical pathways in a stratified environment, therefore the bacteria aligned in the Earth’s field reduce a three dimensional search to a single dimension, swimming up-downwards the stratified column.

Another possible role of the magnetosomes has been suggested as detoxifying agents scavenging reactive oxygen species^6^ or providing resistance to UV-B irradiation^7^. These works suggest that the synthesis of magnetosomes could respond to a survival mechanism against a stressful situation.

In this line, the aim of this work is to analyse the role of the magnetosomes in the tolerance of magnetotactic bacteria to the following transition metals: Co, Mn, Ni, Zn and Cu. All these metals are essential micronutrients for the cell^8^ but can also be toxic at high concentrations. The mechanisms of metal intoxication include oxidative stress, inhibition of the electron transport chain, antagonizing metal uptake and enzyme mismetallation^9,10^. The potential deleterious effects rely on the bioavailability of the metal as well as on different environmental factors (i.e. pH, temperature, organic matter content)^11,12^. In order to deal with such stress, microorganisms have developed several strategies, including the formation of protein-metal complexes to sequester metal ions, the reduction of these to less harmful forms or the continuous pumping of metal out of the cell^10,13^. In the same way, the synthesis of magnetosomes could have evolved as an additional strategy to enhance the tolerance to metals in MTB.

The atomic numbers of Co, Mn, Ni, Zn, Cu are close to Fe, which could facilitate their incorporation into the magnetite structure of the magnetosome mineral core. Magnetite (Fe_3_O_4_) has an inverse spinel structure, in which one third of the Fe atoms occupy tetrahedral positions in the form of Fe^3+^, while the other two thirds occupy octahedral positions, half of them in the form of Fe^2+^ and the other half as Fe^3+^. Metallic cations can be incorporated into the magnetite spinel structure by replacing Fe cations and form metal ferrites, whose general formula is M_x_Fe_3-x_O_4_ (M = metallic cation).

In this work, we investigate the tolerance of the magnetotactic bacterium *Magnetospirillum gryphiswaldense* to metal stress. Firstly, by measuring the sensitivity curves of bacterial populations with and without magnetosomes, we prove that bacteria with magnetosomes have more tolerance to metal stress than bacteria without magnetosomes. Afterwards, we analyze the role of the magnetosomes in the resistance mechanisms of the bacteria against metal stress by means of x-ray absorption near edge structure (XANES) experiments. XANES measurements on both the whole bacteria and on magnetosomes isolated from the bacteria have allowed us to determine whether bacteria store the excess metal in the magnetosomes or in other cell compartments. Our findings reveal that the tolerance mechanisms are metal-specific. While Co is incorporated only in the magnetosomes, Ni is incorporated in cell compartments other than the magnetosomes. Mn, Zn and Cu are found both in the magnetosomes and in other cell compartments. Interestingly, in all cases, the metal content in the magnetosomes is low (< 2 at.%/Fe), and, except for Co, most metal is incorporated in other cell compartments.

## Results and discussion

### Metal sensitivity curves and TEM

Figure 1 shows the sensitivity curves of *M. gryphiswaldense* to Co, Mn, Ni, Zn and Cu with and without magnetosomes. In all cases, the population with magnetosomes shows less sensitivity to metals than the population without magnetosomes, that is, the concentration of metal needed to produce growth inhibition is always higher in the population with magnetosomes, indicating that magnetosomes confer tolerance to metals.

**Figure 1:**
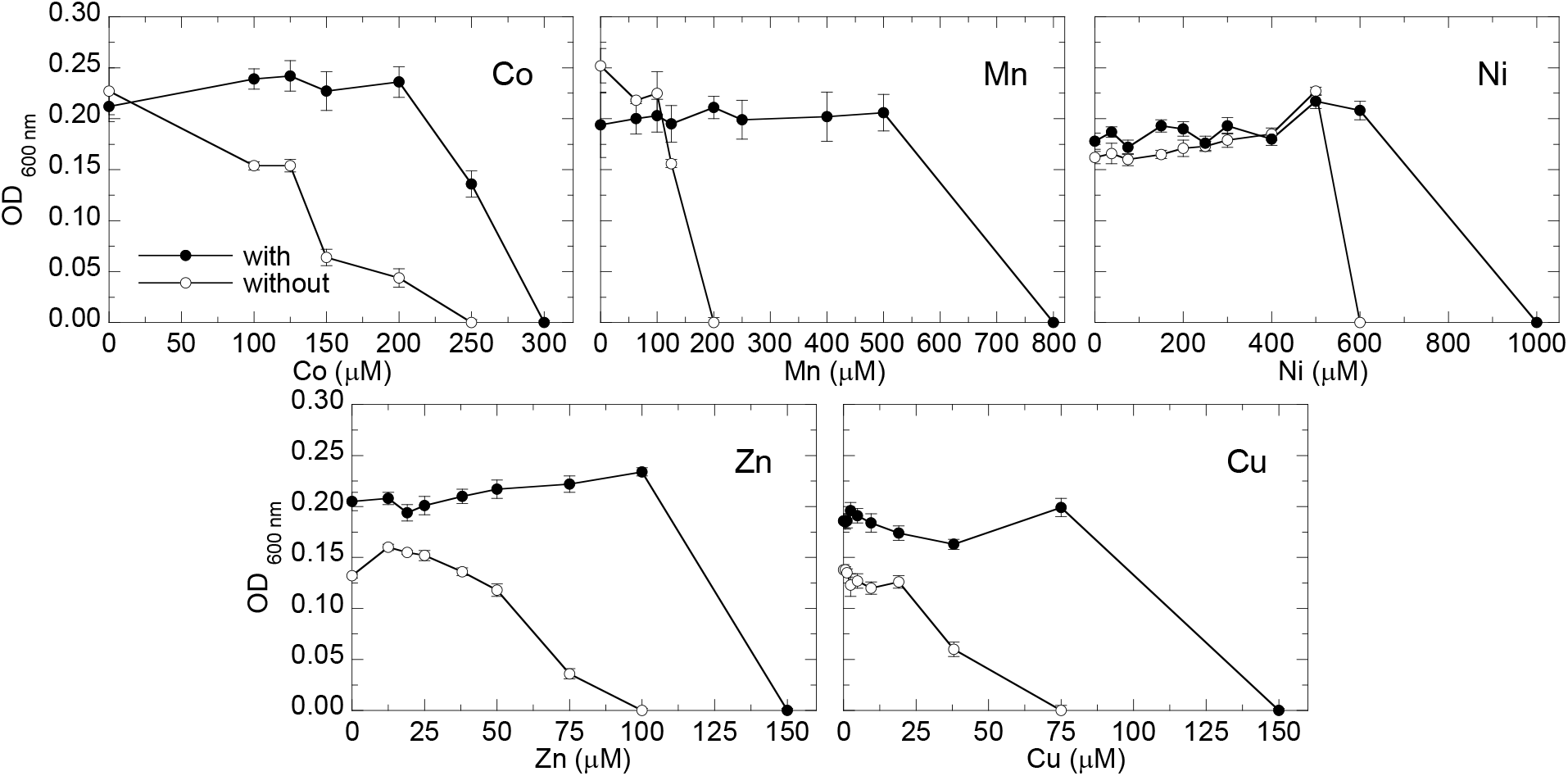
Sensitivity curves (optical density (OD_600_) as a function of metal concentration) to Co, Mn, Ni, Zn and Cu of *M. gryphiswaldense* populations with and without magnetosomes. The mean values and standard deviation (SD) of eight replicates are represented.

Even though to our knowledge no study has been published regarding the role of magnetosomes in the tolerance to metals, there are several studies related with the sensitivity of magnetotactic bacteria to certain metals. Tanaka et al.^14^ determined the minimum inhibitory concentration (MIC) values of the same five metals of this work (Co, Mn, Ni, Zn and Cu) in magnetosomes-bearing *Magnetospirillum magneticum* AMB-1. They reported MIC values for Mn (1000*μ*M) and Zn (200*μ*M) similar to the concentrations that arrest the growth in our case. However, MIC values reported in [14] for Co (60*μ*M), Ni (100*μ*M) and Cu (40*μ*M) still support the growth of magnetosomes-bearing *M. gryphiswaldense*. These different results suggest the development of different strategies against metal stress in both *Magnetospirillum* species.

In order to check possible alterations in magnetosome size and/or arrangement inside the cell due to the presence of metals in the culture medium, a TEM analysis was performed. Figure 2 shows representative TEM images of *M. gryphiswaldense* cultured in a medium supplemented with Co, Mn, Ni, Zn and Cu together with control bacteria cultured in a non-supplemented medium. The images do not show substantial differences among the bacteria in terms of cell morphology, and, as the control, all bacteria have a single magnetosome chain.

**Figure 2:**
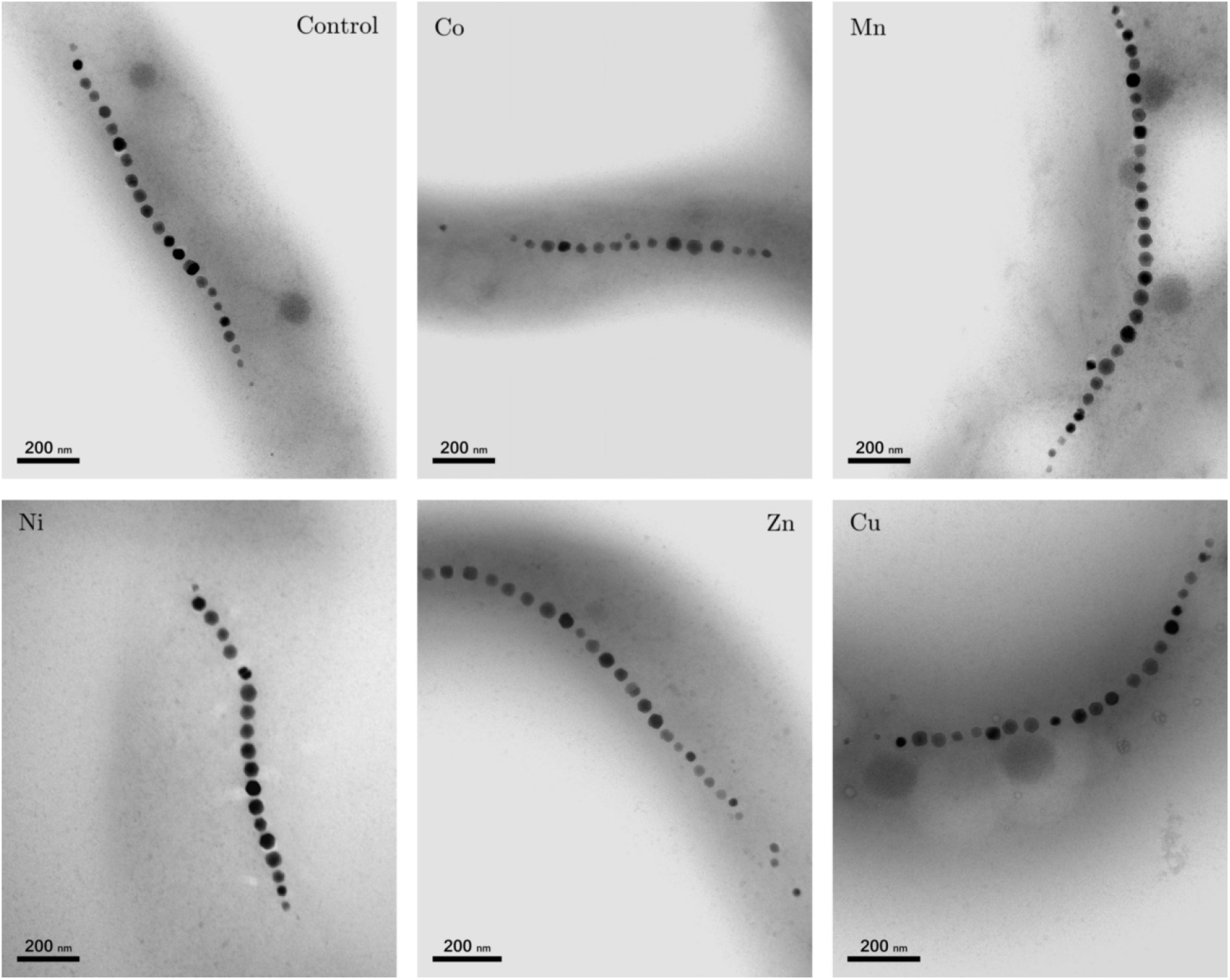
Representative TEM images of control *M. gryphiswaldense* bacteria and bacteria grown in a medium supplemented with Co, Mn, Ni, Zn and Cu. The corresponding magnetosome size distribution and number of magnetosomes per chain are shown in Figure 3.

Figure 3A shows the particle size distributions and the number of magnetosomes per chain for the control cells and the cells cultured in metal-supplemented medium obtained from the TEM imaging. No significant changes in magnetosome size are observed among the magnetosomes synthesized in a metal-supplemented medium and the control. The mean values vary between 34 ± 8 nm for Cu and 37 ± 10 nm for Mn, being the control 35 ± 8 nm. On the contrary, Prozorov et al.^15^ obtained nanoparticles with smaller sizes than the control when culturing bacteria in a Zn enriched medium (32 ± 13 nm in Zn vs. 45 ± 13 nm the control). In *Magnetospirillum magneticum*, Tanaka et al.^14^, did also report a decrease in particle size in the presence of Co, Ni and Cu but no changes in the presence of Mn. However, in *M. magnetotacticum*, Kundu et al.^16^ reported larger sizes in magnetosomes synthesized in the presence of Zn and Ni than in the control (23 ± 3 nm and 25 ± 5 nm for Zn and Ni respectively, and 15 ± 5 nm for the control).

**Figure 3:**
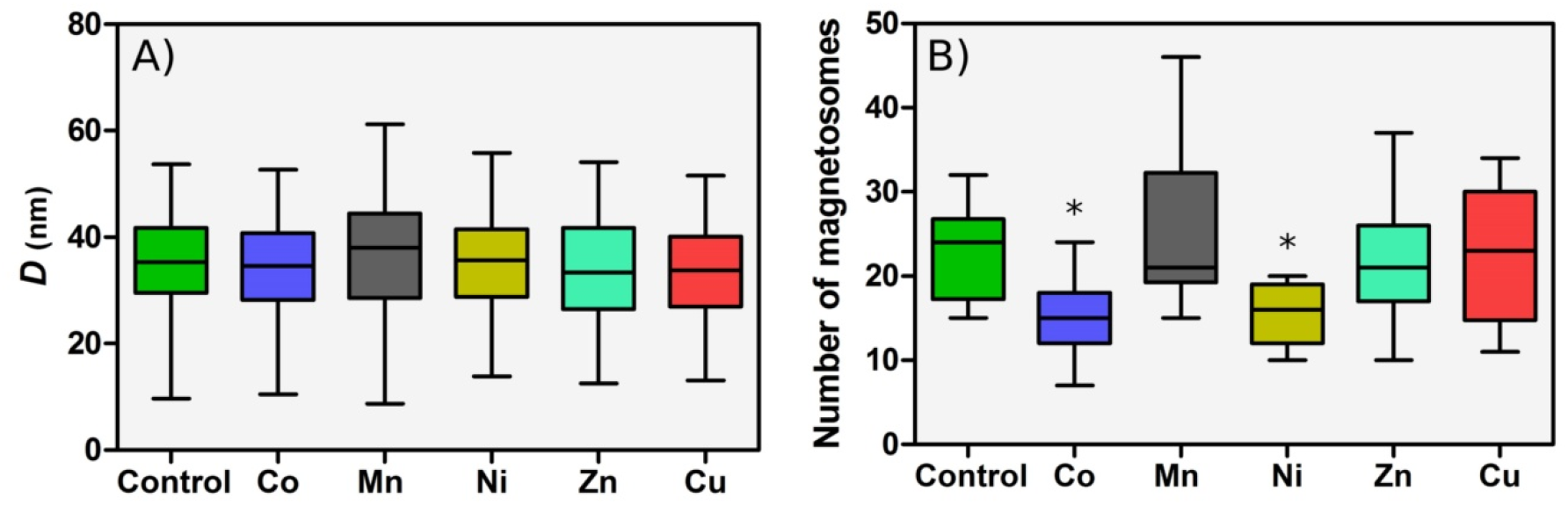
Box plots of magnetosome size (A) and number of magnetosomes per chain (B) of control bacteria and bacteria grown in a metal supplemented medium. Each analysis was performed over approximately 15 bacteria and 250 magnetosomes. The length of each box represents the central range covering 50% of the distribution, the horizontal line represents the median and the bar ends correspond to the maximum and minimum values of each distribution. (* p< 0.05)

Regarding the length of the chains (Figure 3B), bacteria grown with Co and Ni supplements show fewer magnetosomes per chain than the control (16 ± 5 and 15 ± 4 magnetosomes per chain for Co and Ni, respectively, and 23 ± 6 for the control). A similar result was found for *M. magneticum*^14^. No significant differences with respect to the control are observed for bacteria grown with Mn, Zn and Cu supplements. While unusually long chains have been found in the case of Mn, the differences in the mean values with respect to the control are not statistically significant.

### Study of the metal incorporation by XANES spectroscopy

Once confirmed that magnetosomes confer tolerance to metal excess in the growth medium, here we have analyzed whether magnetosomes bearing bacteria do actually incorporate the excess metal in the magnetosomes or whether they incorporate it in any other cell compartment. This analysis has been performed by means of x-ray absorption spectroscopy (XAS). In particular, we analysed the near-edge region of the Fe and metal (M) K-edge XAS spectra (the XANES region). XANES is an element-specific technique that has allowed us to detect the presence of the metal in the sample and to gain information on the oxidation state, coordination chemistry and local symmetry around the absorbing element^17^.

In this work, XANES measurements have been performed on both the whole bacteria and the corresponding isolated magnetosomes. In this way, we have been able to assess whether the metal is incorporated by the bacteria, and if it does, whether it enters the magnetosome. We have also been able to gain information on the average oxidation state and symmetry of the metal sites, and in some cases we have been able to identify the chemical complex the metals are forming by means of a fingerprinting analysis (comparison with XANES spectra of model compounds) and/or linear combination fit (LCF) with model compounds. In the latter case, when more than one compound is contributing to the XANES signal, the LCF allows quantifying the atomic fraction of each contributing phase.

Figure 4 shows the Fe K-edge XANES spectra of the whole bacteria (control and M-bacteria) and the corresponding isolated magnetosomes (control magnetosomes and M-magnetosomes). Regardless of the metal studied, all M-bacterial spectra and the spectra of their corresponding isolated M-magnetosomes are coincident with the control, which corresponds to the spectrum measured for magnetite^18,19^. The spectra of isolated M-magnetosomes being identical to magnetite regardless of the metal studied suggests that the amount of metal incorporated into the magnetosome is low, insufficient to change appreciably the coordination environment of the Fe atoms. In fact, no trace or a very small amount of the M-metal was detected in previous energy-dispersive x-ray spectroscopy (EDS) measurements on isolated M-magnetosomes (see the supplementary information). Finally, the coincidence of the bacterial spectra with the corresponding magnetosomes indicates that, regardless of the low mass fraction of magnetosomes in the bacterial samples (less than 2% of dry weight), the contribution of any other Fe compounds present in the bacteria is masked by the magnetosomes, indicating that most of the Fe in the bacteria is in the magnetosomes.

**Figure 4:**
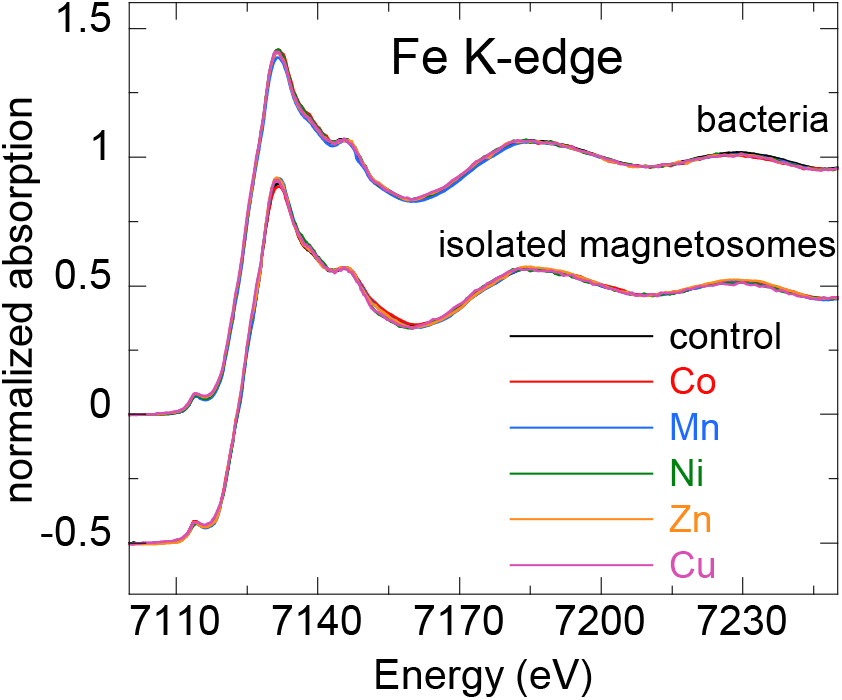
Normalized Fe K-edge XANES spectra of control bacteria and bacteria grown in a metal supplemented medium (M-bacteria), and magnetosomes isolated from the control bacteria and from M-bacteria (M-magnetosomes). The latter have been shifted to ease comparison.

Figures 5 to 9 show the normalized M K-edge XANES spectra of the control bacteria (bacteria grown in a metal non-supplemented medium), M-bacteria and corresponding isolated M-magnetosomes. Each metal will be analyzed separately in the following.

**Figure 5:**
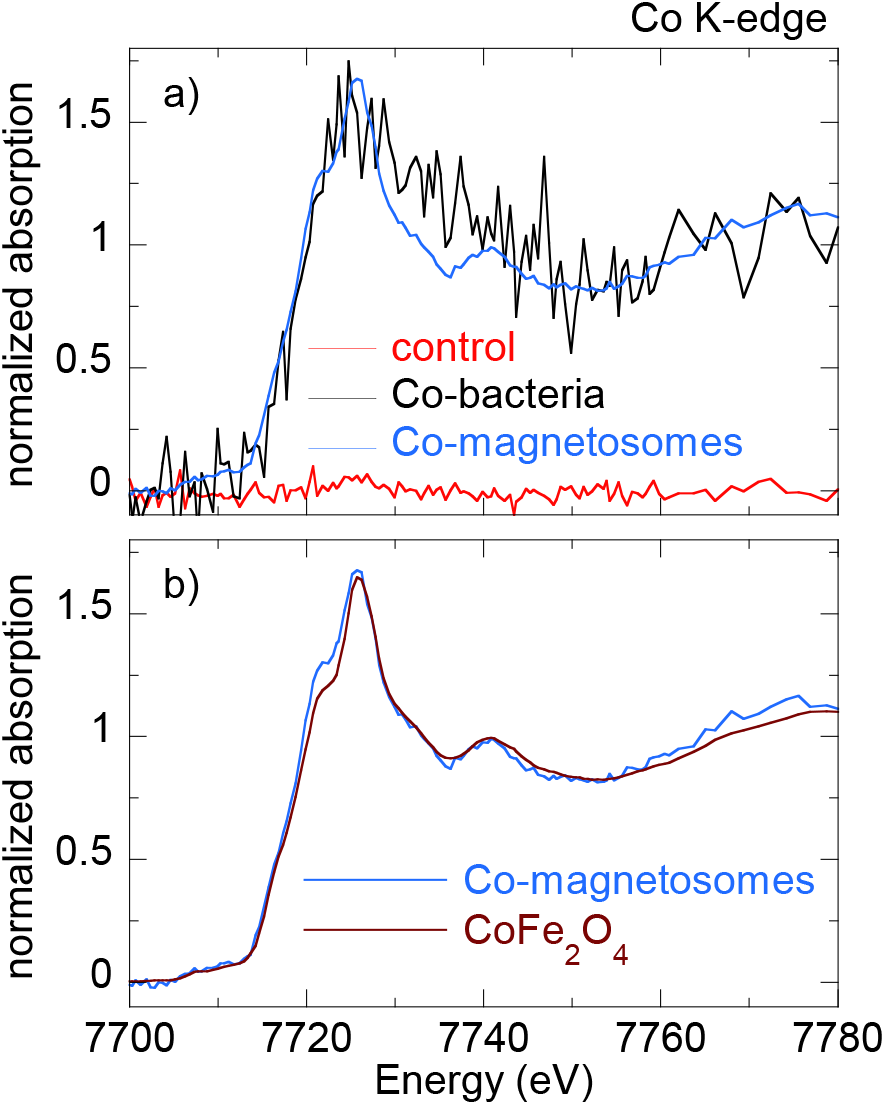
a) Normalized Co K-edge XANES spectra of control bacteria, bacteria grown in a Co supplemented medium (Co-bacteria) and magnetosomes isolated from Co-bacteria (Co-magnetosomes). b) Co K-edge XANES spectrum of Co-magnetosomes compared to Co ferrite (CoFe_2_O_4_).

#### Co

As shown in Figure 5a, regardless of Co being an essential micronutrient, there is no detectable Co absorption in control bacteria. However, a measurable although noisy Co K-edge XANES spectrum is obtained for Co-bacteria, meaning that the incorporation of Co is promoted when bacteria are grown in a Co-supplemented medium.

The isolated Co-magnetosomes show a well-defined Co K-edge XANES spectrum that confirms that Co is incorporated into the magnetosome structure. The presence of Co in the magnetosomes suggests that the poor signal obtained in Co-bacteria could be entirely coming from the Co-magnetosomes, indicating that most Co is stored in the magnetosomes and not in any other cell compartment.

The spectrum of the Co-magnetosomes matches with the one of Co ferrite (CoFe_2_O_4_) (Figure 5b), indicating that Co is introduced into the spinel structure of the magnetite and, as in CoFe_2_O_4_, it occupies octahedral sites in the Co^2+^ oxidation state. Note that, according to EDS (see the supplementary information), the Co content in the magnetosomes is only 1.6% at./Fe, thus the Co ferrite in the magnetosomes is not stoichiometric (Co_0.05_Fe_2.95_O_4_). In spite of the low Co content, it has been previously reported that the magnetic properties of Co-magnetosomes change notably with respect to the undoped ones^20–22^.

#### Mn

The control bacteria show a well defined Mn K-edge XANES spectrum, suggesting that Mn is an abundant element in the cell in basal conditions (Figure 6a). As shown in Figure 6b, comparison of the energy edge position of this spectrum with Mn references (MnO_2_ (Mn^4+^), Mn_2_O_3_ (Mn^3+^), MnO (Mn^2+^)) suggests that the cell mainly incorporates the Mn in Mn^2+^ compounds. Additional references have been included in the supplementary information (Figure S3).

**Figure 6:**
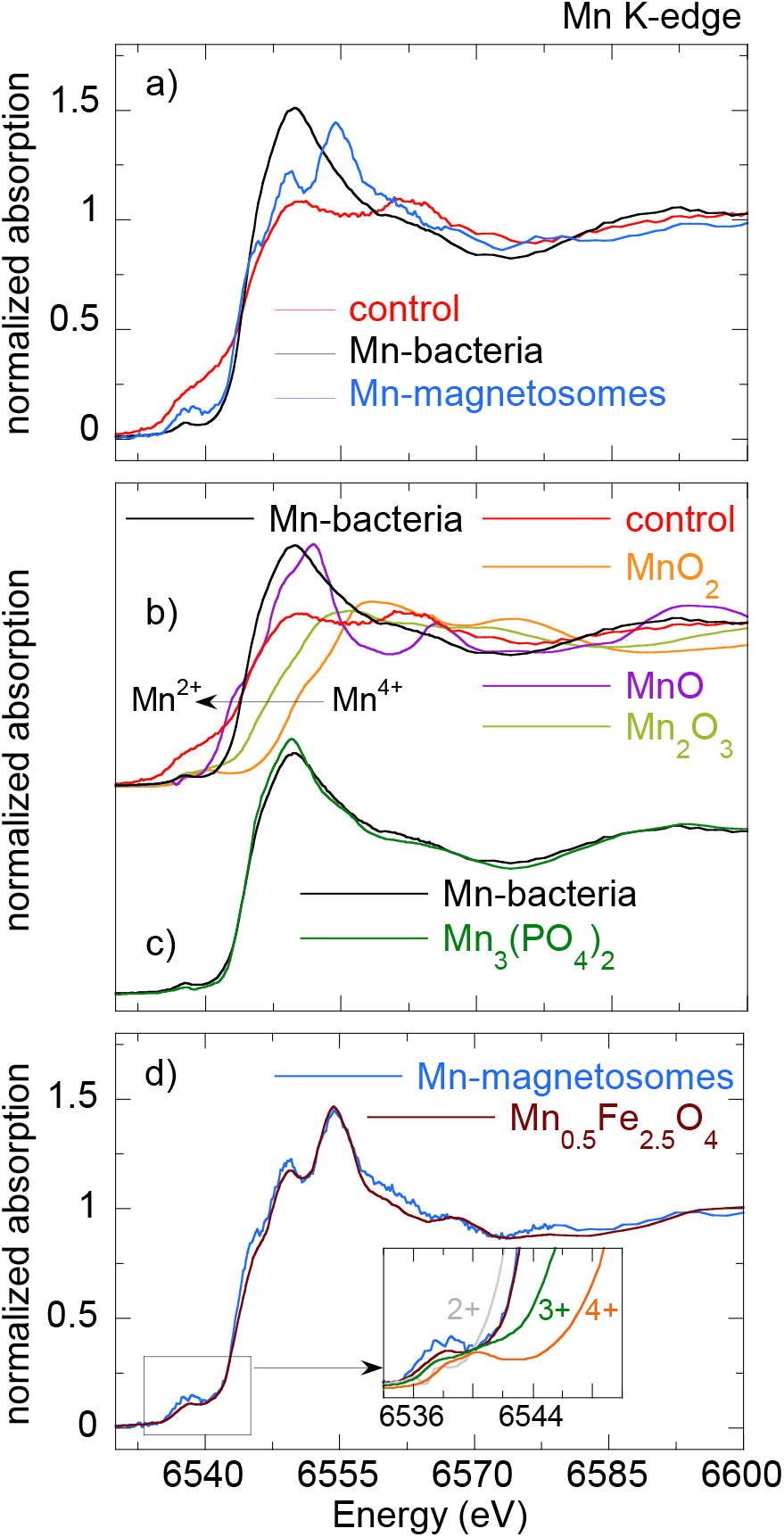
a) Normalized Mn K-edge XANES spectra of control bacteria, bacteria grown in a Mn supplemented medium (Mn-bacteria) and magnetosomes isolated from Mn-bacteria (Mn-magnetosomes). b) Mn K-edge normalized XANES spectra of Mn-bacteria and the control together with Mn references: MnO_2_ (Mn^4+^), Mn_2_O_3_ (Mn^3+^), MnO (Mn^2+^). c) Comparison of the XANES spectra of Mn-bacteria and Mn_3_(PO_4_)_2_ and d) Mn-magnetosomes and Mn_0.5_Fe_2.5_O_4_ ^23^. The inset in d) highlights the edge position of the Mn-magnetosomes with respect to the references.

The XANES spectrum of the Mn-bacteria differs from the control, which means that an excess of Mn promotes a change in the average coordination environment of Mn ions. This suggests that bacteria incorporate the Mn in different cell compartments depending on the concentration of Mn in the medium, although comparison to the references suggests that Mn retains its Mn^2+^ oxidation state (Figure 6b). The similarity of the spectrum of Mn-bacteria with Mn^2+^ phosphate (Mn_3_(PO_4_)_2_) (Figure 6c) suggests that bacteria could be incorporating the excess Mn in the polyphosphate granules present in the cytoplasm of *gryphiswaldense*, supporting previous works that indicate that magnetotactic bacteria incorporate Mn and Zn in phosphorous-rich cell inclusions^24^.

The isolated magnetosomes present a well-defined XANES spectrum that confirms that Mn is also incorporated into the magnetosome. The Mn phase in the isolated Mn-magnetosomes can be identified as a Mn ferrite (Mn_x_Fe_3-x_O_4_) from direct comparison with Mn_0.5_Fe_2.5_O_4_ ^23^ (Figure 6d). The oxidation state of Mn in this phase can be inferred from the position of the absorption edge. Indeed, as shown in the inset of Figure 6d, the absorption edge of Mn-magnetosomes is between MnO (Mn^2+^) and Mn_2_O_3_ (Mn^3+^), approximately 1 eV above the absorption edge of Mn^2+^. Knowing that there is a linear relationship between the edge position and the oxidation state^25^, and since there is a 3.6 eV shift from Mn^2+^ to Mn^3+^, a 1 eV shift indicates that Mn in Mn-magnetosomes holds a mixed valence with 75% Mn^2+^ and 25% Mn^3+^, in agreement with what is reported for Mn ferrites^26,27^.

Therefore, bacteria incorporate Mn both in the magnetosomes and in other cell compartments, particularly in the polyphosphate granules. However, in the Mn-bacterial XANES spectrum (Figure 6c), which should include the contribution of the Mn in the polyphosphate granules and in the Mn-magnetosomes, the latter contribution is negligible, suggesting that most Mn is incorporated into the polyphosphate granules rather than in the magnetosomes.

#### Ni

As shown in Figure 7a, the Ni K-edge XANES spectrum of the control sample displays a poor signal-to-noise ratio, coherent with a low Ni content. The XANES signal improves considerably in Ni-bacteria, confirming that bacteria incorporate Ni when grown in a Ni-supplemented medium. According to the edge position, Ni is in the Ni^2+^ oxidation state in the bacteria (Figure 7b) but this spectrum could not be identified with the available standards.

**Figure 7:**
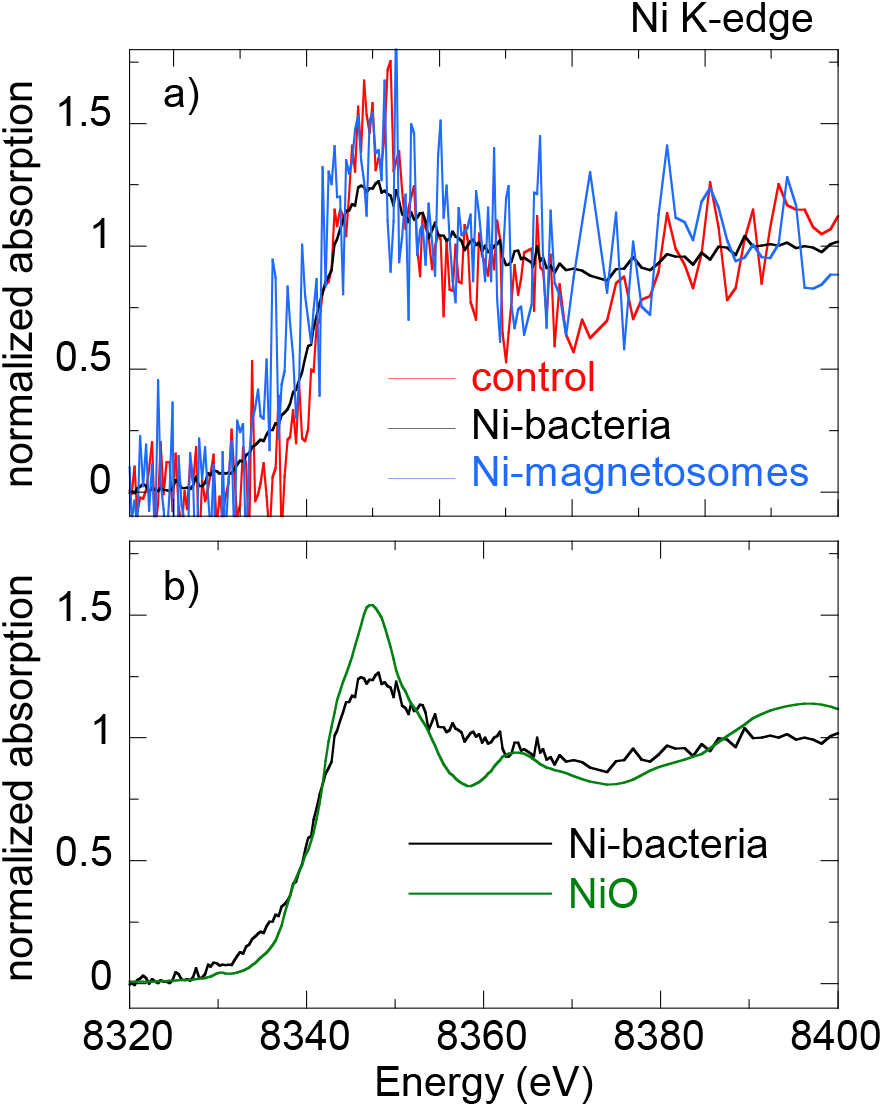
a) Normalized Ni K-edge XANES spectra of control bacteria, bacteria grown in a Ni supplemented medium (Ni-bacteria) and magnetosomes isolated from Ni-bacteria (Ni-magnetosomes). b) Ni K-edge XANES spectra of Ni-bacteria and NiO (Ni^2+^).

Regarding the Ni-magnetosomes, the absorption is extremely small, indicating that Ni is not incorporated in the magnetosome and therefore all the Ni incorporated by Ni-bacteria is in other cell compartments.

#### Zn

The control sample shows a clear Zn K-edge XANES spectrum (Figure 8a), an indication of the cell incorporating Zn in basal conditions.

**Figure 8:**
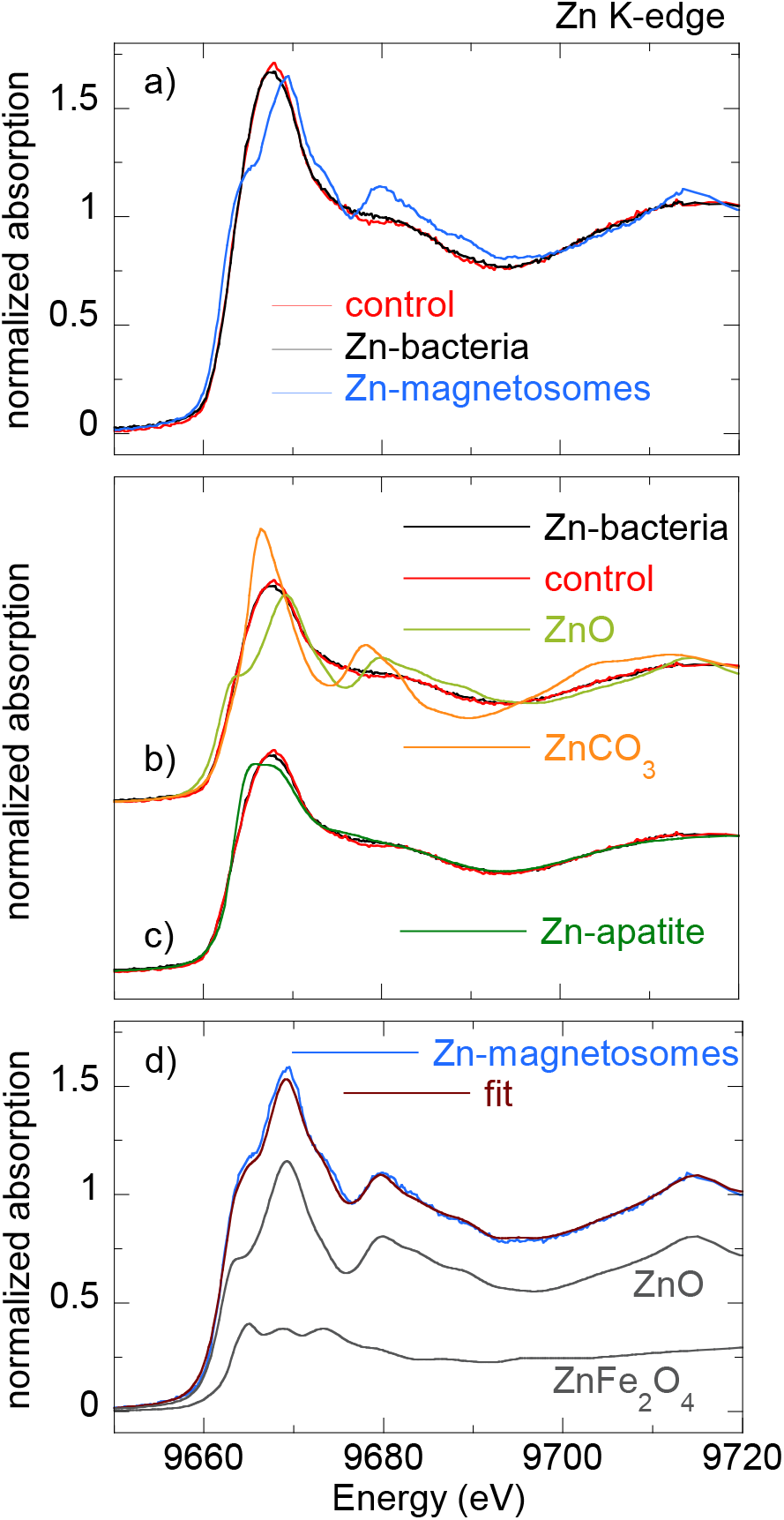
a) Normalized Zn K-edge XANES spectra of control bacteria, bacteria grown in a Zn supplemented medium (Zn-bacteria) and magnetosomes isolated from Zn-bacteria (Zn-magnetosomes). b) Zn K-edge XANES spectra of Zn-bacteria and control together with two Zn^2+^ standards: ZnO and ZnCO_3_. c) Comparison of XANES spectra of Zn-bacteria and Zn- control with Zn apatite. d) Linear combination fit of the Zn K-edge XANES spectrum of Zn- magnetosomes with 72% ZnO and 28% Zn ferrite (ZnFe_2_O_4_).

When the medium is supplemented with Zn, the XANES spectrum of Zn-bacteria is very similar to the control, indicating that Zn is mostly incorporated in the same chemical species as in basal conditions. Figure 8b shows the XANES spectra of Zn-bacteria and control bacteria together with some Zn^2+^ references. Here it is clear that Zn in the bacteria is in the Zn^2+^ oxidation state. However, phase identification by fingerprinting or by a linear combination fit (LCF) of the available standards (see available standards in the supplementary information, Figure S4) has been unsuccessful. As shown in Figure 8c, the closest similarity has been found with Zn adsorbed onto hydroxyapatite, a partially distorted phase with a preferential tetrahedral coordination where oxygen atoms compose the first coordination shell around the Zn^2+^ ions, and phosphorous the second shell^28,29^. The similarity with Zn hydroxyapatite suggests that Zn in the bacteria could be embedded in a similar coordination environment, perhaps in the polyphosphate granules as Mn^24^.

Regarding the isolated Zn-magnetosomes, the Zn K-edge XANES spectrum (Figure 8a) evidences that Zn is incorporated into the magnetosome. A LCF analysis (Figure 8d) suggests that the obtained spectrum results from the coexistence of two phases in which Zn has a tetrahedral coordination: ZnO (72%) and Zn ferrite ZnFe_2_O_4_ (28%). This indicates that some Zn is in fact inserted into the spinel structure of the magnetite forming ZnFe_2_O_4_, but that there are Fe depleted zones in the magnetosomes where Zn ions retain the tetragonal coordination forming ZnO.

Therefore, in a Zn-supplemented medium, bacteria incorporate Zn both into the magnetosome and in other cell compartments, the same cell compartments as in basal conditions. The contribution of the Zn-magnetosomes is masked in the Zn-bacteria XANES spectrum, indicating that most Zn is incorporated in other cell compartments rather than in the magnetosomes.

#### Cu

As shown in Figure 9a, control bacteria have enough Cu in basal conditions to provide a clear XANES spectrum. This spectrum changes slightly when bacteria are grown in a Cu-supplemented medium. Both spectra display a feature at the edge jump that in the Cu-bacteria is more intense and slightly shifted to higher energies. The feature at the edge position is characteristic of Cu^1+^ complexes, and its position and intensity can change as a function of the coordination environment^30^. In fact, comparison with reference standards (Figure 9b) confirms the Cu^1+^ oxidation state. Studies on other bacterial strains do also report that Cu^1+^ is the dominant valency in which Cu binds to the bacteria^31^. As shown in Figure 9c, the similarity between the spectra of Cu-bacteria and covellite (CuS or Cu_3_(S_2_)S, a Cu^1+^ compound with hardly no Cu^2+^ character^32^) supports works that report that Cu^1+^ largely bounds sulphydryl group-containing ligands as metallothionein proteins and other low molecular mass thiols^31,33,34^ and suggests that an additional Cu phase is promoted in the cell when bacteria are cultured in a Cu-rich medium.

**Figure 9:**
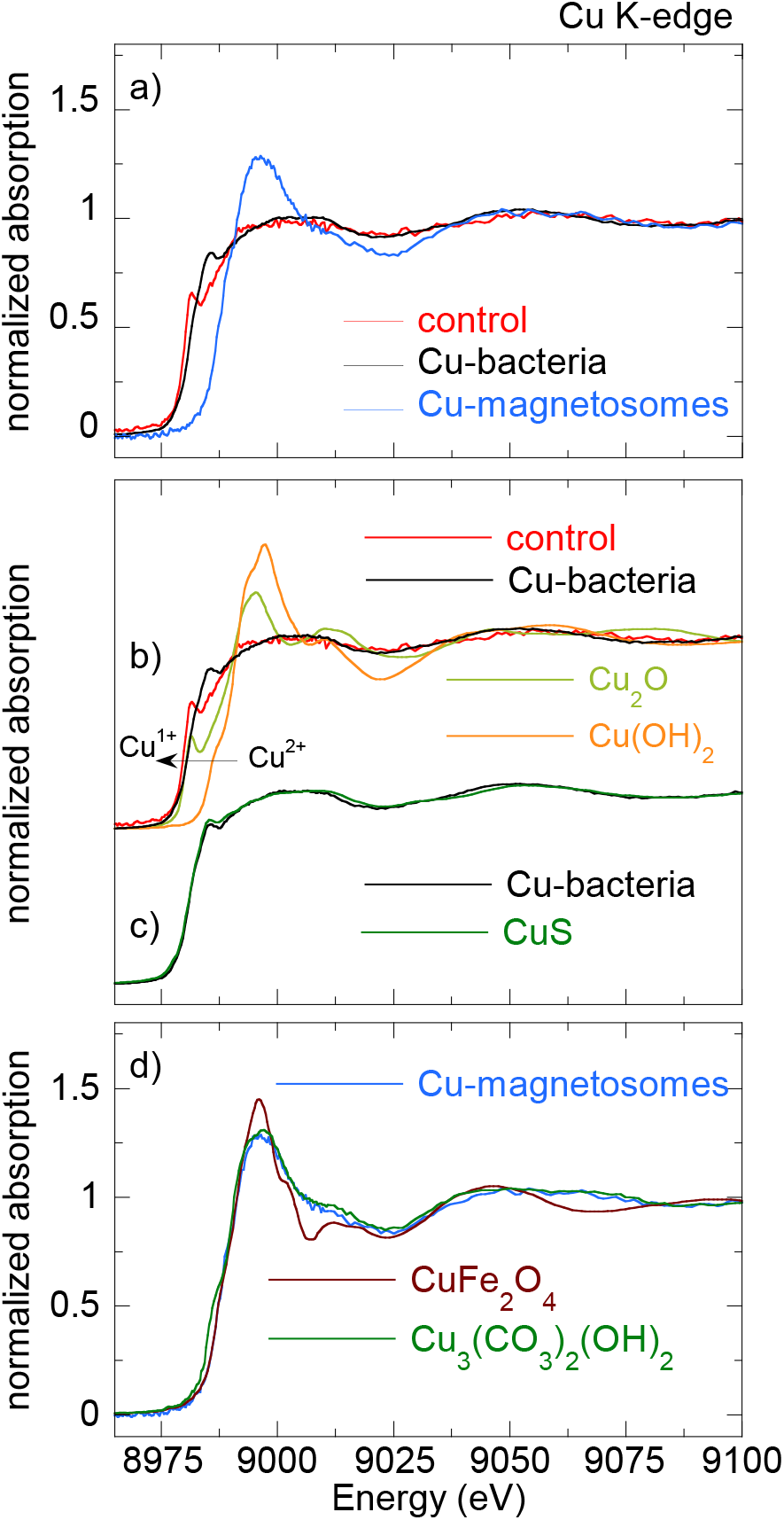
a) Normalized Cu K-edge XANES spectra of control bacteria, bacteria grown in a Cu supplemented medium (Cu-bacteria) and magnetosomes isolated from Cu-bacteria (Cu-magnetosomes). b) Cu K-edge XANES spectra of Cu-bacteria and control together with two Cu standards: Cu_2_O (Cu^1+^) and Cu(OH)_2_ (Cu^2+^). c) Cu-bacteria compared with CuS. d) Cu-magnetosomes compared with Cu ferrite (CuFe_2_O_4_) and azurite (Cu_3_(CO_3_)_2_(OH)_2_).

On the other hand, the isolated Cu-magnetosomes display a well-defined XANES signal, which confirms that Cu is in the magnetosomes. The energy edge is shifted to higher energies with respect to the bacterial energy edges, indicating that Cu in the magnetosomes is in a more oxidized state. As shown in Figure 9d, the Cu K-edge spectrum of the isolated magnetosomes does not match the one of Cu ferrite (CuFe_2_O_4_), indicating that Cu does not enter the magnetite mineral structure. However, the coincidence of the energy edge position confirms that Cu is in a Cu^2+^ oxidation state in the magnetosome. A good match is obtained when comparing Cu-magnetosomes with the Cu compound azurite (Cu_3_(CO_3_)_2_(OH)_2_). This is a carbonate Cu compound, which suggests that Cu could be incorporated in the magnetosome membrane.

Finally, the spectrum of Cu-bacteria cannot be reproduced by means of a LCF including the contributions of the Cu-magnetosomes and any other combination of spectra such as the control, CuS, and/or other available standards, suggesting that most of the Cu incorporated by the bacteria is in cell compartments different from the magnetosomes.

In summary, to cope with the excess of metal in the growth medium, *M. gryphiswaldense* exhibit different mechanisms depending on the metal tested. In this way, only in the case of Co do bacteria incorporate the metal solely in the magnetosomes. On the contrary, Ni is not incorporated in the magnetosomes, and can only be found in other cell compartments. In the case of Mn, Zn and Cu, bacteria do accumulate metal in the magnetosomes, but most of the incorporated metal is stored in other cell compartments. This is evidenced in the M-bacteria XANES spectra, where the contribution of the M-magnetosomes is masked by the metal content of the whole M-bacteria.

In any case, the amount of metal incorporated into the magnetosome structure is always small, thus the tolerance to transition metals exhibited by magnetosome-bearing bacteria can not be explained solely by the metal-sequestering capacity of magnetosomes and other detoxifying mechanisms based on the catalytic power of the magnetosomes could be involved.

## Conclusion

We have investigated the role of magnetosomes in the tolerance of *M. gryphiswaldense* to transition metal stress. The profiles of the sensitivity curves of bacteria with and without magnetosomes grown in metal-supplemented media show that the tolerance to Co, Mn, Ni, Zn and Cu increases notably when cells synthesize magnetosomes. These results point out that magnetosomes not only confer a magnetotactic behavior but also could respond to a survival mechanism against a stressful situation. The resistance mechanisms triggered in magnetosome-bearing bacteria under metal stress have been investigated by means of x-ray absorption near edge spectroscopy (XANES). XANES spectroscopy has shown that metal uptake in magnetosome-bearing bacteria is enhanced when culturing in metal enriched media, although the metal is not always introduced into the magnetosome structure. Under metal stress the bacteria incorporate metals either in the magnetosomes or in other cell compartments, depending on the metal. In any case, the metal content incorporated in the magnetosomes is always small. Thus the storage of metals in the magnetosomes cannot fully explain the long survival of magnetosome-bearing bacteria under metal stress, and other detoxifying mechanisms based on the catalytic power of the magnetosomes could be involved. Future research on mutant strains deficient in magnetosome production could shed light on this open question in search of such alternative detoxifying mechanisms based on the presence of magnetosomes.

## Materials and methods

### Bacterial strain and standard growth conditions

The MTB used in this work belong to the species *Magnetospirillum gryphiswaldense* strain MSR-1 (DMSZ 6631) that synthesize cuboctahedral magnetite particles. The strain was cultured in the flask standard medium (FSM) described by Heyen & Schüler^35^ (per litre of deionized water: 0.1 g KH_2_PO_4_, 0.15 g MgSO_4_7H_2_O, 2.38 g Hepes, 0.34 g NaNO_3_, 0.1 g yeast extract, 3 g soy bean peptone). The medium contained 0.3% (wt/vol) sodium pyruvate as a carbon source and Fe(III)-citrate was added at a final concentration of 100 *μ*M. Incubation proceeded for 96 h at 28°C without agitation in loosely stoppered bottles with a headspace-to-liquid ratio of approximately 1:4 and air was used in the headspace. To obtain bacteria without magnetosomes the medium was not supplemented with Fe(III)-citrate (low iron medium, LIM) and incubation proceeded with shaking (120 r.p.m.). To discard any effect of the iron content in the culture medium on the bacterial growth, the generation time was calculated for bacteria with and without magnetosomes cultured in both FSM and LIM. The generation time ranges between 7.0 and 7.6 hours and no significant differences were found among sets (see Table S1).

### Metal sensitivity curves

The sensitivity analysis to metals (M=Co, Mn, Ni, Cu, Zn) was performed on bacterial populations with and without magnetosomes. Serial dilutions of every metal in culture medium were prepared in 96-well microplates. For the sensitivity curves of magnetosomes bearing bacteria, the metal dilutions were prepared in FSM. In the case of bacteria without magnetosomes, metal dilutions were made in LIM. The dilution range tested was specific for each metal: Co (0 − 600*μ*M), Mn (0 − 1000*μ*M), Ni (0 − 1000*μ*M), Zn (0 − 200*μ*M) and Cu (0 − 150*μ*M). The metals were added in the form of organic or inorganic M^2+^ salts: Co (C_12_H_10_Co_3_O_14_) and Mn (C_12_H_10_Mn_3_O_14_) citrates, and Ni (NiSO_4_), Zn (ZnSO_4_) and Cu (CuSO_4_) sulfates. Dilutions were inoculated with bacteria with or without magnetosomes in stationary phase to achieve an initial concentration of 10^6^ cell mL^−1^. After an incubation period of 120 hours at 28°C under slight shaking, the optical density (OD_600_) of the culture was measured to evaluate bacterial growth. Eight replicates were performed for each metal by inoculating from the same bacterial culture.

Selected dilutions showing bacterial growth were analyzed by electron microscopy (TEM). The analysis was performed on unstained cells adsorbed onto 300 mesh carbon-coated copper grids. TEM images were obtained with a Philips CM120 electron microscope at an accelerating voltage of 100 kV. The particle size distribution was analyzed using a standard software for digital electron microscope image processing, ImageJ^36^.

### X-ray absorption near edge spectroscopy (XANES)

XANES measurements were performed on the Fe and M (M=Co, Mn, Ni, Zn, Cu) K-edges on bacteria with magnetosomes (control bacteria and bacteria grown in a metal supplemented medium, M-bacteria) and on isolated magnetosomes (M-magnetosomes).

Bacteria were grown up to stationary phase in FSM supplemented with: Co (100*μ*M); Mn (100*μ*M); Ni (100*μ*M); Zn (50*μ*M); Cu (10*μ*M). Cells were collected by centrifugation (8000*g* for 15 minutes at 4°C). The cell-free supernatants were decanted, acidified with nitric acid to reach pH 2-3 and analyzed by inductively coupled plasma atomic emission spectroscopy (ICP-AES) to determine the remaining amount of M-metal after bacterial growth. Analyses were performed three times. From the ICP-AES results the M-metal uptake by the bacteria was estimated around 20-30% of the supplied M-metal. The obtained pellets were fixed in 2% glutaraldehyde, washed three times with 10 mM HEPES (pH 7.4), freeze dried and compacted in 5 mm diameter pills. Magnetosomes were isolated according to the protocol described by Muela et al.^37^. Briefly, cells were disrupted by French press and the magnetosomes were isolated from the lysate using a magnetic rack and rinsed 10 times with 10 mM HEPES-200 mM NaCl (pH 7.4). The purified magnetosomes were freeze-dried, mixed with sacarose and compacted in 5 mm diameter pills.

XANES measurements were performed at the BL22 CLÆSS beamline of the ALBA synchrotron facility (Barcelona, Spain) and at the branch A of the BM25 (SpLine) beamline of the ESRF synchrotron facility (Grenoble, France)^38^. Both experiments were performed at room temperature. The monochromator used in the experiments was a double Si crystal oriented in the (111) direction.

Fe K-edge measurements were performed in transmission mode, as well as the M K-edge measurements of commercial reference samples (CoFe_2_O_3_, MnFe_2_O_4_, NiFe_2_O_4_, ZnFe_2_O_4_, CuFe_2_O_4_, Mn_3_(PO_4_)_2_, MnO, MnO_2_, Mn_2_O_3_, C_12_H_10_Mn_3_O_14_, MnATP, MnEDTA, NiSO_4_, ZnSO_4_, Zn acetate, CuO, Cu_2_O, CuOH_2_, CuSO_4_, Cu acetate, CuS, Cu_3_(CO_3_)_2_(OH)_2_). In addition, the Mn K-edge XANES reference Mn_0.5_Fe_2.5_O_4_ was provided by Dr Mazarío^23^, and Zn K-edge XANES references (ZnO, ZnSO_2_, smithsonite (ZnCO_3_), Zn adsorbed onto hydroxyapatite, Zn_3_(PO_4_)_2_, Zn-cysteine, Zn-histidine, Zn-glutathione, Zn-proline, hemimorphite (Zn_4_(Si_2_O_7_)(OH)_2_H_2_O)) were provided by Dr. Meneghini, from Università di Roma Tre, Italy^28^.

Due to the low M-metal concentration of the samples, the M K-edge measurements of the bacterial and isolated magnetosome samples were performed in fluorescence yield mode using a silicon-drift diode (SDD) solid-state detector (ALBA) or an energy-dispersive 13-element Si(Li) multidetector (ESRF). Under these conditions, the detection limit of the technique is as low as 10 ppm (10 mg/kg)^39^. From three to five spectra were acquired for each sample and merged to improve the signal-to-noise ratio. Metal foils were measured in both experiments and used as references to calibrate the energy so that the edge position of the sample can be determined with an accuracy of 0.2 eV.

The experimental spectra were normalized using standard procedures for background subtraction and data normalization as implemented in the free software Athena of the Ifeffit package^40,41^.

## Acknowledgments

The Spanish and Basque Governments are acknowledged for funding under projects number MAT2017-83631-C3-R and IT-1245-19, respectively. Dr. L. Marcano acknowledges the financial support provided through a postdoctoral fellowship from the Basque Government. The authors acknowledge the technical and human support provided by SGIker (UPV/EHU). The ESRF, MCIU, and CSIC are acknowledged for the provision of ESRF synchrotron radiation facilities. The authors thank Dr. Carlo Meneghini for providing the XANES spectra of Zn standards and R. Fernández-Pacheco for his assistance in the EDS measurements.

## Notes

### Competing Interest Statement

The authors have declared no competing interest.

