## Supplementary material for "Magnetosomes could be protective shields against metal stress in magnetotactic bacteria"

June 2, 2020

### 1 Generation time

Table S1 shows the growth parameters of *M. gryphiswaldense* with ($Mag^+$) and without ($Mag^-$) magnetosomes in a medium supplemented with Fe (FSM) and in a medium not supplemented with Fe (LIM). The generation time ranges between 7.0 and 7.6 hours. The populations analyzed in the article correspond to ($Mag^+$, FSM) and ($Mag^-$, LIM).

| **Inoculum** | **Culture medium** | $\mu$ **(h$^{-1}$)** | **g (h)** |
|---|---|---|---|
| $Mag^+$ | FSM | 0.094 $\pm 0.002$ | 7.4 $\pm 0.2$ |
| $Mag^+$ | LIM | 0.092 $\pm 0.004$ | 7.6 $\pm 0.3$ |
| $Mag^-$ | FSM | 0.099 $\pm 0.004$ | 7.0 $\pm 0.3$ |
| $Mag^-$ | LIM | 0.095 $\pm 0.003$ | 7.3 $\pm 0.2$ |

Table S1: Specific growth constant ($\mu$) and generation time ($g$) of bacterial populations with ($Mag^+$) and without ($Mag^-$) magnetosomes in a medium supplemented with Fe (FSM) and in a medium not supplemented with Fe (LIM). Each value is the mean value and the standard deviation (± SD) of five replicates. The populations analyzed in the article correspond to ($Mag^+$, FSM) and ($Mag^-$, LIM).

#### 2 Energy-dispersive x-ray spectroscopy (EDS) on isolated M-magnetosomes

To estimate quantitatively the amount of M-metal incorporated into the M-magnetosomes isolated from the bacteria we have carried out a chemical analysis by means of energy-dispersive x-ray spectroscopy (EDS) in TEM mode.

EDS was performed on extracted magnetosomes adsorbed either onto 300 mesh carbon-coated copper grids (for Mn, Co and Ni-magnetosomes) or onto 300 mesh lacey-carbon molybdenum grids (for Cu and Zn-magnetosomes), to avoid the superposition of the absorption lines of the grid with the energies corresponding to the metal of interest. Images were obtained with a Philips CM200 electron microscope at an accelerating voltage of 200 kV which includes an EDS detector. EDS spectra were acquired with a counting time of 5 min to optimize a good signal-to-noise ratio and to minimize the induced radiation damage. The beam size was 65 nm, allowing the microanalysis of small clusters of several magnetosomes. Thus, EDS was performed on selected areas, some of which are represented by white squares in the TEM-micrographs displayed on the left panel of Figure S1. To prevent radiation damage, the selected areas were in different regions of the sample.

The right panel of Figure S1 displays the EDS spectra obtained from different regions of M-magnetosome samples. All the spectra show two main peaks. The one at lower energies ($\approx 6400$ eV) corresponds to the Fe-K$_\alpha$ emission line, and the one at higher energies ($\approx 7060$ eV), to the Fe-K$_\beta$. Mn, Co and Ni samples present also two intense peaks at 8038 and 8905 eV, which correspond to Cu emission energies arising from the copper grid. The dashed lines mark the M-K$_\alpha$ emission lines.

Except for Co-magnetosomes, the presence of metal M in the isolated M-magnetosomes is undetectable by this technique, which means that the M content is below the resolution limit,

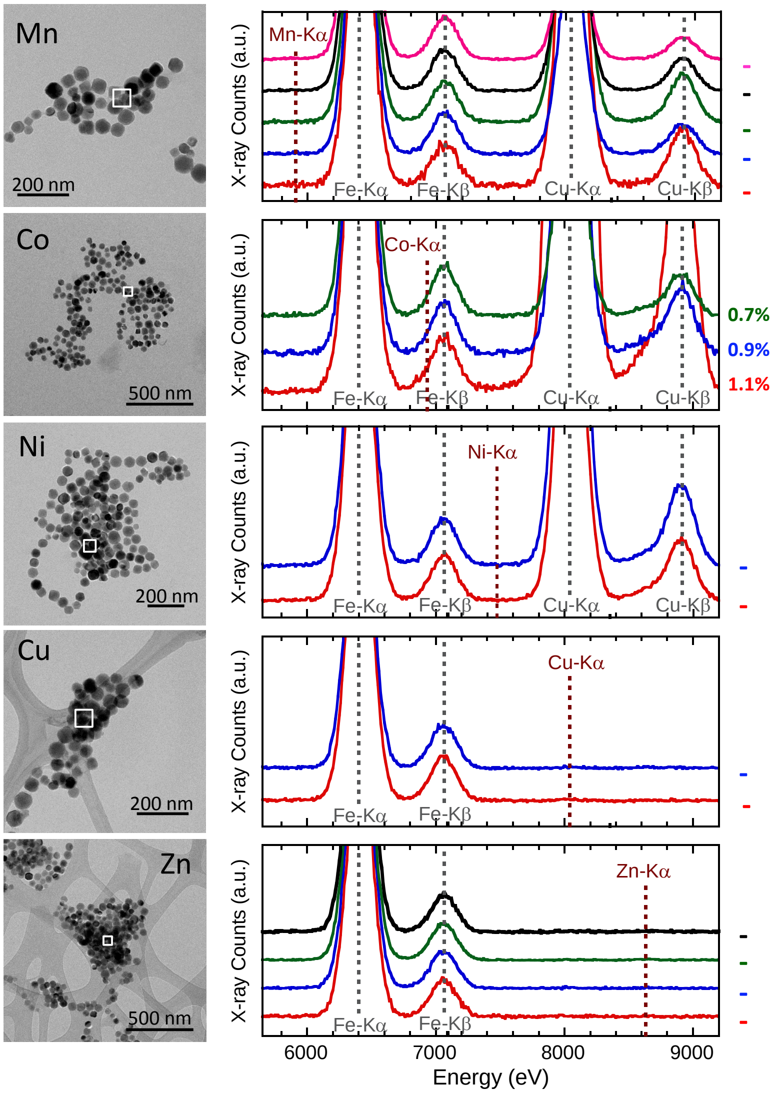


Figure S1: EDS analysis carried out on small clusters of isolated M-magnetosomes. Left panel: TEM micrographs of isolated M-magnetosomes. The white squares delimit some of the regions analyzed by EDS (beam size 65 nm). Right panel: EDS spectra acquired on the selected regions. Dashed lines indicate the position of M-K$\alpha$ emission lines.

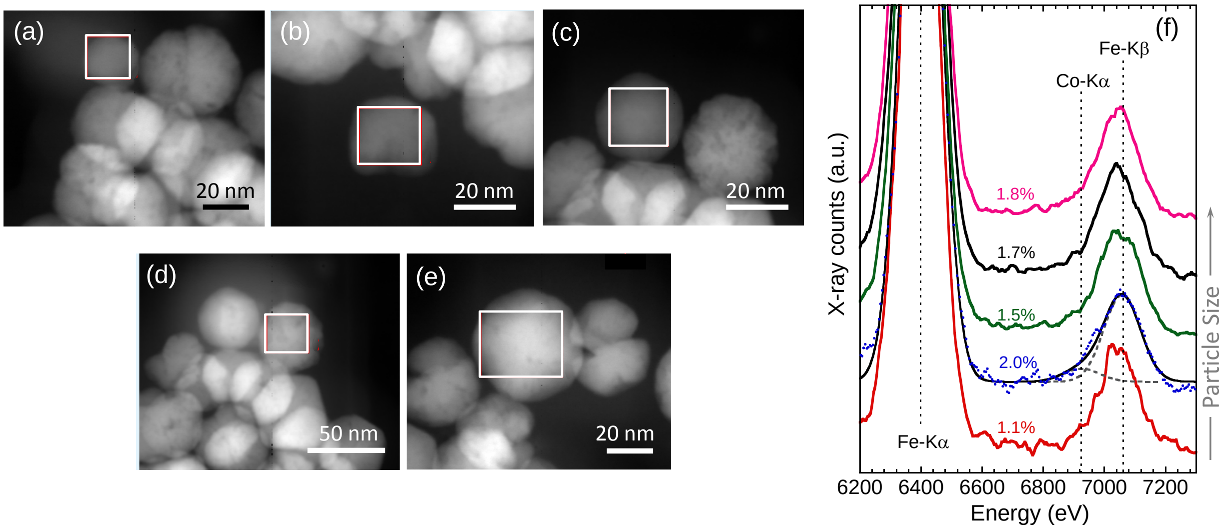


Figure S2: STEM-HAADF images of individual Co-doped magnetosomes with increasing particle size: a) 25, b) 28, c) 40, d) 43 and e) 45 nm. White squares represent the regions analyzed by EDS. f) EDS spectra of the five individual Co-magnetosomes. Values linked to each spectrum point out the Co at.% resulting from the Gaussian fits. An example of the Gaussian fit is shown for the 28 nm-sized magnetosome. Dashed lines mark the x-ray emission lines of Fe and Co.

which is around 1-2 at.%.

In the case of Co, the Co-$K_\alpha$ emission line (at $\approx 6920$ eV) introduces a slight asymmetry in the low energy region close to the Fe-$K_\beta$ peak. By means of a Gaussian fit of the Fe-$K_\alpha$, Fe-$K_\beta$ and Co-$K_\alpha$ emission lines, the Co content, estimated as the ratio of the integrated areas of the Fe-$K_\alpha$ and Co-$K_\alpha$ peaks (Co/(Co+Fe)), is 0.9 at.% Co. Since this value is in the order of magnitude of the EDS resolution limit, to obtain the Co at.% more accurately, an additional EDS analysis was performed in an equipment with a higher resolution.

In this way, isolated Co-magnetosomes were analyzed by Scanning-Transmission Electron Microscopy (STEM) with a High Angle Annular Dark Field Detector (HAADF, Fischione) in a FEI Tecnai F30 electron microscope at an accelerating voltage of 200 kV (Instituto de Nanociencia de Aragón). In this mode, the intensity is proportional to the thickness and the square of the atomic number, therefore chemical elements with a higher atomic number appear with a higher intensity. Figures S2a-e show the STEM-HAADF images of isolated Co-magnetosomes. The white squares display the regions on which each EDS spectrum was obtained (EDAX detector, 0.5 eV/pixel). The sample was scanned on the displayed squared regions, where the EDS signals were accumulated.

Five different magnetosomes with increasing size, from 25 nm (a) to 48 nm (e), were analyzed. The resulting EDS spectra are shown in Figure S2f. A Gaussian fit analysis (an example is shown for the 28 nm sized particle) shows that the Co content ranges between 1.1 and 2% of the total Co and Fe atoms and is not correlated to the particle size, giving a mean Co content of 1.6 at.% ($\pm 0.5\%$).

These results are in agreement with values reported in the bibliography for *M. gryphiswaldense* doped magnetosomes: Prozorov et al.[1] report a 1% Mn content in Mn-doped magnetosomes, and Staniland et al.[2] report 1.4% in Co-doped magnetosomes. In contrast, higher metal contents have been found in *M. magneticum* of 15.6%, 3% and 2.7% in Cu, Mn and Cu-doped magnetosomes, respectively, although no Zn and Ni was detected in Zn,Ni-doped magnetosomes[3].

#### 3 Mn K-edge XANES references

Figure S3 shows the Mn K-edge XANES spectra of the references used in this work together with the Mn-bacteria and the control.

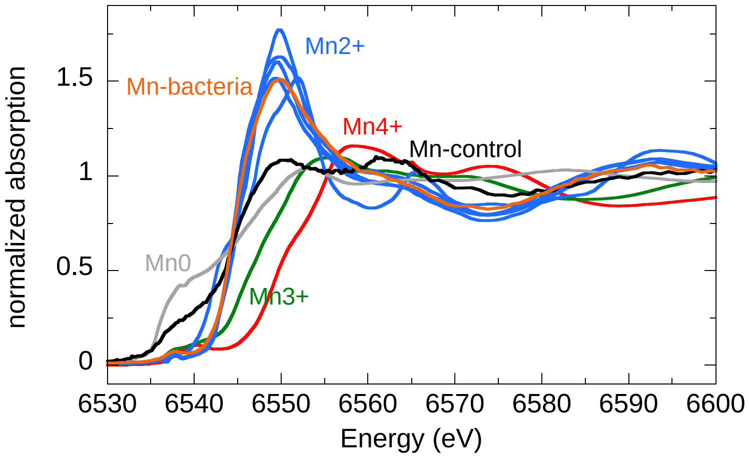


Figure S3: Mn K-edge XANES spectra of Mn-bacteria and the control together with some Mn references.

### 4 Zn K-edge XANES references

Figure S4 shows the Zn K-edge XANES spectra of the references used in this work together with the Mn-bacteria and the control.

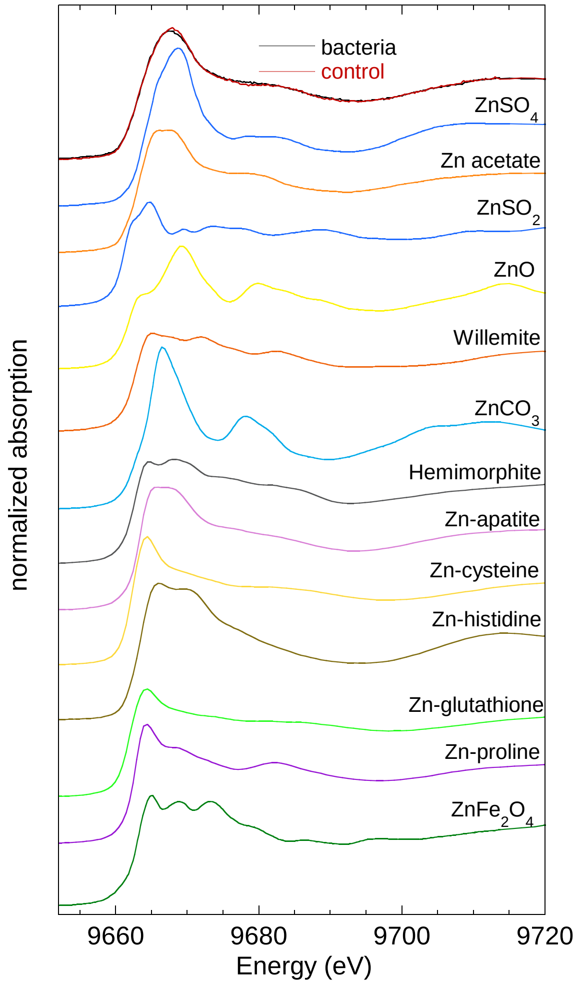


Figure S4: Zn K-edge XANES spectra of Zn-bacteria and the control together with the available Zn references. All the references except for Zn acetate, $ZnSO_4$ and Zn ferrite ($ZnFe_2O_4$) were provided by the group of Dr C. Meneghini, from the Università di Roma Tre, and some of them can be found in reference 4.
